# Heavy-tailed prior distributions for sequence count data: removing the noise and preserving large differences

**DOI:** 10.1101/303255

**Authors:** Anqi Zhu, Joseph G. Ibrahim, Michael I. Love

## Abstract

In RNA-seq differential expression analysis, investigators aim to detect genes with changes in expression across conditions, despite technical and biological variability. A common task is to accurately estimate the effect size. When the counts are low or highly variable, the simple effect size estimate has high variance, leading to poor ranking of genes by effect size. Here we propose *apeglm*, which uses a heavy-tailed Cauchy prior distribution for effect sizes, resulting in lower bias than previous shrinkage estimators, while still reducing variance. *apeglm* is available at http://bioconductor.org/packages/apeglm, and can be used from within the *DESeq2* software.

## Background

RNA sequencing (RNA-seq) is a widely used assay for measuring the expression of transcripts from the genome. One common goal is to identify which genes are differentially expressed (DE) between experimental conditions, and to estimate the strength of the difference. The difference is usually defined in terms of the logarithmic fold change (LFC) between average expression levels of different conditions. The expression level of a gene in an RNA-Seq experiment is proportional across samples to a scaled count, representing the number of observed single-or paired-end reads that could be assigned to a given gene at a given library size. Scaling for the library size of the experiment is necessary, and other scaling factors can be included as well [1–3]. Many variations on the standard RNA-seq protocol exist, as well as other sequencing-based assays such as chromatin immunoprecipitation followed by sequencing (ChIP-seq), and to the degree that these other experiments assess differences in scaled counts using estimated LFCs, the methods described here are generally applicable to these other assays as well.

Many statistical methods have been developed for differential expression analysis of RNA-seq [4–13]. Their common approach in detecting differentially expressed (DE) genes is to find sets of genes such that the null hypothesis of no difference in expression between conditions can be rejected, usually targeting the false discovery rate (FDR) for the set. However, a gene can be found significantly different, and the null rejected, even if the size of difference is very small [13]. For further research interests, rather than only considering the order of genes according to adjusted or unadjusted *p*-values, it is also of interest to order genes by the estimated effect size itself (the LFC).

It is challenging to accurately estimate the LFCs for genes with low expression levels, or genes with a high coefficient of variation. Due to experimental costs and time, RNA-seq experiments designed for hypothesis generation typically have a small number of biological replicates (n of 3 — 5) for each condition group [7]. When the counts of sequenced reads are small or have a high coefficient of variation in one or a subset of the conditions, the estimated LFCs will have high variance, leading to some large estimated LFCs, which do not represent true large differences in expression. One approach that reduces the problem of these noisy LFC estimates is to filter out low count genes. The authors of *edgeR* [7] and *limma-voom* [12] suggest using a filtering rule that removes genes with low scaled counts before statistical analysis [14]. Other methods take a Bayesian modeling approach, including *ShrinkBayes* [9] and *DESeq2* [13]. *DESeq2* applies an adaptive *Normally*-distributed prior, to produce a shrinkage estimator for the LFC for each gene. However, in our analysis, we found that filtering or application of Normal priors each can have drawbacks, either leading to loss of genes with sufficient signal, or overly aggressive shrinkage of true, large LFCs.

In this article, we present an empirical Bayes procedure that stabilizes the estimator of LFCs, without overly shrinking large LFCs, and uses the posterior distribution for point estimates and posterior probabilities, such as the the aggregated *s*-value [15] and the false-sign-or-smaller rate. We extend the basic framework of *DESeq2*, a Negative Binomial (NB) generalized linear model (GLM) [16] with moderated dispersion parameter, by exchanging the Normal distribution as a prior on LFCs with a heavy-tailed Cauchy distribution (a t distribution with 1 degree of freedom). We use various approximation techniques to provide Approximate Posterior Estimation for the GLM (*apeglm*). We compare *apeglm* to four existing methods on two benchmarking RNA-seq datasets. We demonstrate the advantages of *apeglm*’s shrunken estimates in reducing variance while preserving the true large effect sizes. We also show that *apeglm* shrunken estimates improve gene rankings by LFCs, relative to methods which do not apply Bayesian shrinkage on the LFCs. *apeglm* is available as an open-source R package on Bioconductor, and can be easily called from within the *DESeq2* software.

## Results

### Strong filtering thresholds may result in loss of DE genes

It is difficult to accurately estimate the LFCs for genes with low read count; maximum likelihood estimates (MLE) of LFCs for genes with low read count have high variance due to the dominance of sampling variance over any detectable biological differences. The MLEs of LFCs for these genes may not reflect the true biological difference of gene expression between conditions, and thus are not reliable for plotting or ranking genes by effect size [13]. Chen et al. [14] suggested to remove from analysis the genes that have low scaled counts across samples. They define a scaled quantity, the *counts per million* (CPM), which is the counts *Y_gi_* divided by a robust estimator for the library size, multiplied by one million. The filtering rule is to keep only those genes with *n* or more samples with CPM greater than the CPM value for a raw count of 10 for the least sequenced sample. The suggested value for *n* is the sample size of the smallest group. CPM filtering occurs prior to any statistical analysis. Other data-independent thresholds, such as requiring a CPM of 0.5 or 1 from *n* or more samples can be even more aggressive at removing genes with potential signal when the sequencing depth is high.

We illustrate how filtering can lead to loss of DE genes using the dataset by Bottomly et al. [17], which contains 10 and 11 samples of RNA-seq data for mouses from two strains, C57BL/6J(B6) and DBA/2J(D2), respectively. We repeatedly randomly picked three samples from each strain, balancing across the three experimental batches. We then applied a CPM filtering rule to each random subset, repeating the process 100 times. For all genes in the full dataset, we used *DESeq2* [13] to test for differential expression across strains controlling for batch, defining a set of genes with a nominal FDR threshold of 5%. Supplementary Figure 1 shows four example genes that were removed more than 50% of the time across random subsets, but were reported as differentially expressed by *DESeq2* on the full dataset. There were 207 such genes, which are shown in Supplementary Figure 2. These genes did have information to contribute: for example, they had on average the same sign of estimated LFCs 99% of the time when comparing to the LFCs from the full dataset. These genes, despite having low gene expression, may still be biologically relevant, so we considered statistical methods that produce LFC estimates with low variance for relatively low count genes as well. To be clear, we do not argue against *any filtering*, only against strong filtering for the purposes of obtaining precise LFCs which may discard genes with a relevant signal.

Besides filtering, an additional approach, besides filtering, to produce precise effect sizes is to use scaled pseudocounts, or *prior counts*, to obtain shrinkage estimates of LFCs. The prior count approach is employed by *edgeR* and *limma-voom.* However, setting a prior count does not make use of the statistical information contained in the data for estimating the LFCs, such that the optimal prior count needs to be adapted per dataset. For example, as the sample size increases, the optimal prior count should go to zero, and so a fixed prior count may be sub-optimal. Furthermore, the prior count approach, while helping with high LFC variance from genes with low counts, helps less for high variance genes. Finally, we note that the prior count approach does not provide a posterior distribution for effect sizes, which may be useful for certain analyses discussed below.

### Overview of the *apeglm* method

Following the basic framework of generalized linear models, we propose an adaptive Bayesian shrinkage estimator (Figure 1). We employed a heavy-tailed prior distribution on the effect sizes, where the shape of the prior distribution is fixed, and the scale is adapted to the distribution of observed MLE of effect size for all genes (see Methods). For each gene, the method uses a Laplace approximation to provide the mode of the posterior distribution as a shrinkage estimate, the posterior standard deviation as a measure of uncertainty, and posterior probabilities of interest described below. Our method obviates the need for filtering rules or prior counts, and takes advantage of the statistical information in the data for estimating the effect size. The method is general for various likelihoods, but here we apply it to RNA-seq using a Negative Binomial GLM, where the effect size is a particular LFC (log fold change between groups or an interaction term in a complex design). For genes that have low counts or high variance, this method shrinks the LFCs towards zero thus alleviating the problem of unreliably large LFC estimates.

**Figure 1:**
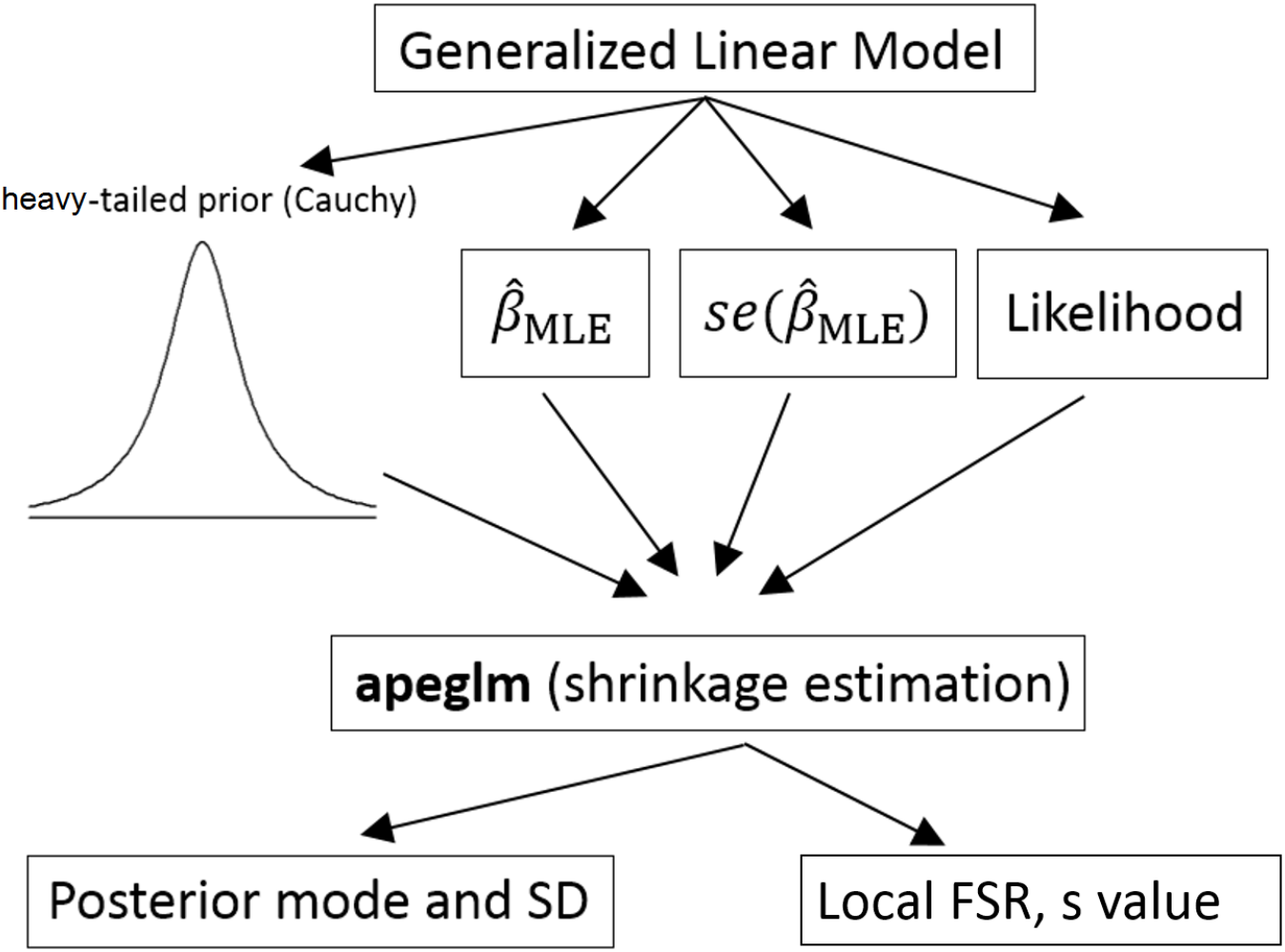
An overview of the method. *apeglm* takes the MLE estimates and corresponding standard deviations (SDs) of a GLM model as input. In *apeglm* we provide a heavy-tailed prior distribution on the coefficients, and computed the shrinkage estimators and corresponding SDs. Users can also define a likelihood function that describes the data and feed to *apeglm. apeglm* also provides the local false sign rates (FSR) and *s*-values [15] as part of the output.

The local false sign rate [15] (FSR) is defined as the posterior probability for a gene that the sign of the estimated effect size is wrong. Similar to the false sign rate, we also make use of a local false-sign-or-smaller *(FSOS*) rate: the posterior probability of having mis-estimated the sign of an effect size, *or the effect size being smaller than a pre-specified value.* For the FSR and FSOS rates, *apeglm* provides an aggregate quantity, the *s*-value proposed by Stephens [15], which can be used for generating lists of genes. The *s*-value for a gene is defined as the average of local FSR over the set of genes that have smaller local FSR than this one (likewise for FSOS, see Methods).

### An adaptive prior controls the false sign rate

We performed an initial assessment of our approach on simulated data, to confirm that the adaptive prior would control the aggregate false sign rate (FSR), when thresholding on s-values, for datasets with varying spread of true LFCs. Using a *fixed*, non-adaptive prior scale leads to loss of control of FSR when the true LFCs were drawn from a Normal distribution with small variance (Supplementary Figure 3). In contrast, matching the scale of the prior to the scale of the true distribution of LFCs regained control of FSR (Supplementary Figure 3). While a prior *smaller* in scale than the true distribution of LFCs also controlled the FSR, it lead to an increase in the relative error of point estimates (Supplementary Figures 4 and 5). Therefore we chose to set the scale of the prior to the estimated scale of the true LFCs using the MLEs and their standard error (Methods).

### Evaluation on highly replicated yeast dataset

To investigate the precision of various estimates of LFCs, we used a highly replicated RNA-seq dataset designed for benchmarking [19]. This dataset consists of RNA-seq data of *Saccharomyces cerevisiae* from two experimental conditions: 42 replicates in *wild-type* (WT) and 44 replicates in a Δ*snf2* mutant. We randomly picked 3 samples from each experimental condition to form a test dataset, and applied differential gene expression methods to estimate the LFCs. We compared the LFCs estimates against the log_2_ ratio of mean scaled counts in the full dataset, which was taken as a “gold standard” LFCs. We repeated the random sampling 100 times. We also performed this same experiment using a sample size of 5 vs 5. For this evaluation and all others, we minimized the influence of genes with no signal for estimating the LFCs, by only evaluating the methods over genes with an average of more than one scaled count per sample. This minimal filtering does not advantage *apeglm.*

We compared the performance of *apeglm* with four other methods for estimation of effect size in RNA-seq, *DESeq2, edgeR, limma-voom*, as well as *ashr* [15]. *ashr* provides generic methods for adaptive shrinkage estimation, taking as input a vector of estimated *β_g_*, i.e. 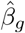, and the corresponding estimates of standard errors. For *ashr*, we input 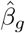 and corresponding standard error using the MLE from *DESeq2* (“*ashr DESeq2* input”), and the estimated coefficient from *limma-voom*, plus a standard error calculated using the moderated variance estimate (“*ashr limma* input”). We also included *edgeR* with a prior count of 5, which helps to moderate the variance of the estimated LFCs from genes with low counts, (*edgeR-PC5*).

Stratifying genes by the absolute value of true LFCs allows us to see where the different methods excel and fall short systematically, across 100 iterations of sub-sampling. *limma* and *edgeR* had the lowest mean absolute error (MAE), with *DESeq2* having the highest error for the largest bin of true LFCs (Figure 2a and c), as was expected from this method. The other shrinkage estimators *apeglm* and *ashr* (with either input) maintained a middle range of MAE. *edgeR-PC5* had low error for the small true LFCs, but then increased to higher error for the largest bin of true LFCs, especially when the sample size increased to 5 vs 5, where the bias approaches that of *DESeq2*.

**Figure 2:**
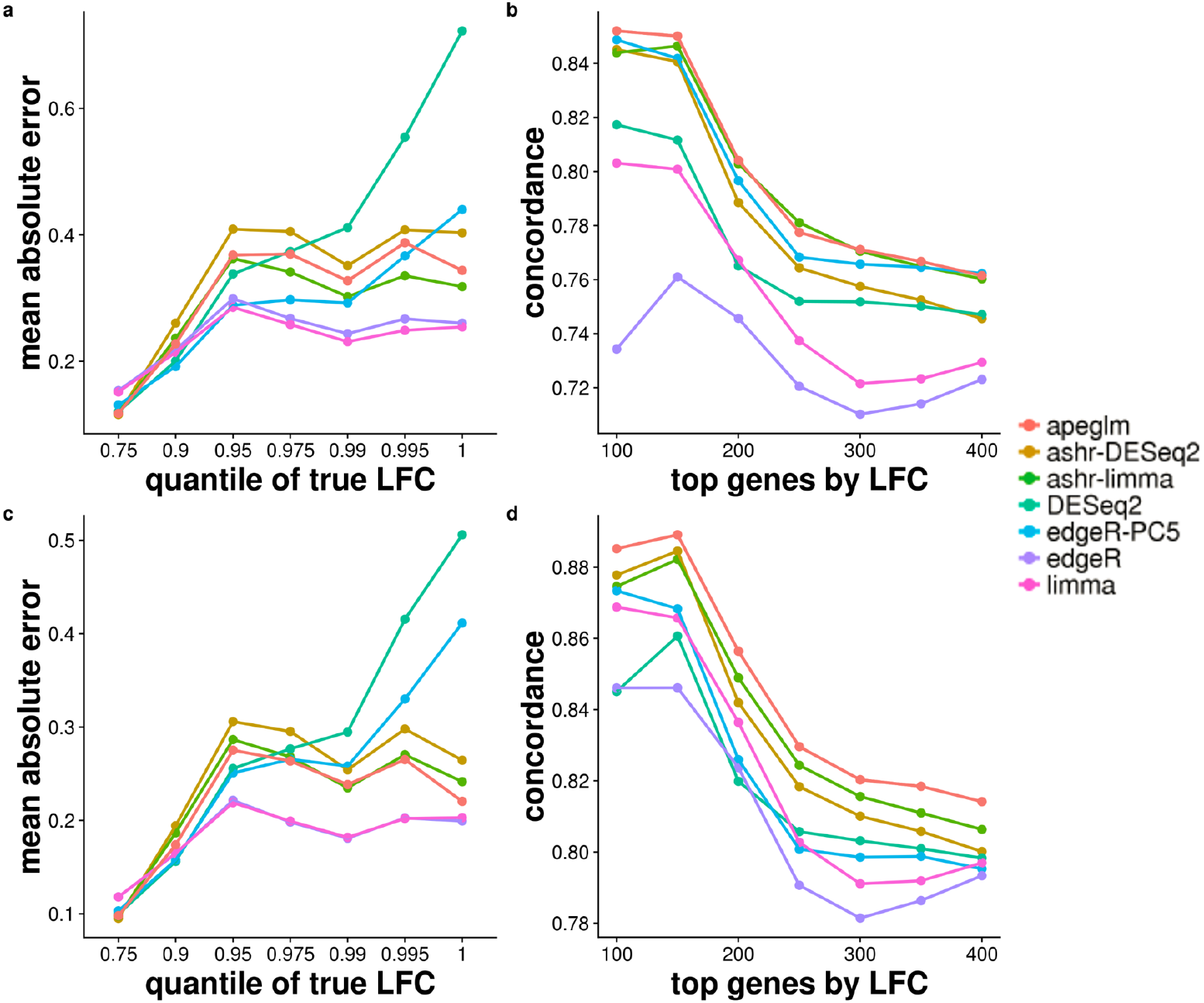
(a) Mean of Absolute Error (MAE) of estimates for 3 vs 3 samples, defined as the mean of the absolute value of the differences between the estimated and reference LFCs, stratified by absolute value of reference LFCs. The mean of MAE over 100 iterations is plotted for each method. The x-axis label gives the upper bound of the bin on absolute value of LFCs. (b) Concordance At the Top (CAT) plot [18] comparing ranked gene lists from each method against the reference ranked gene list for 3 vs 3 samples. Number of top genes ranked by the absolute value of the LFCs is on the x-axis, and the proportion of concordance between the two rankings is on the y-axis. For example, if the ranked gene list from *apeglm* estimated and reference LFCs share 85 of top 100 genes, then the *apeglm* point would fall at (100, 0.85). (c) MAE plot of estimates for 5 vs 5 samples. (d) CAT plot for 5 vs 5 samples.

Ranking genes by estimated LFCs can assist with further investigation into the genes most affected in their expression by changes in condition. We compared the concordance of the top ranked genes by absolute LFC estimates (Figure 2b and d). We examined, for the top *w* genes ranked by absolute value of estimated LFCs, the proportion which were among the top *w* genes by absolute value of reference LFCs (*w* ∈ {100,150,200,…, 400}). *apeglm, ashr* (with either input), and *edgeR-PC5* had the highest concordance of top ranked genes by absolute LFC overall, for 3 vs 3. *apeglm* and *ashr* (with either input) had the highest concordance for the 5 vs 5 sub-sampling experiment. *limma* and *edgeR* tended to have lower concordance compared to shrinkage estimators. *DESeq2* had relatively low concordance among the shrinkage estimators for the smallest *w*. Considering both the MAE stratified by LFCs and the concordance results (Figure 2), we found *apeglm, ashr* and *edgeR-PC5* strike a good balance in estimating the effect size for the 3 vs 3, and *apeglm* and *ashr* for the 5 vs 5 sub-sampling experiment.

In one iteration of random sampling, much of the behavior that was seen systematically over all iterations can be observed (Supplementary Figure 6). *apeglm, ashr* with *DESeq2* or *limma* input, and *edgeR-PC5* did well in estimating LFCs, with LFC estimates close to reference LFCs for most of the genes. *DESeq2* had similar performance to *apeglm*, but was too aggressive in shrinkage for genes with large reference LFCs. *edgeR* and *limma* returned large estimated LFCs for some genes with reference LFCs around zero, which is problematic for ranking genes by effect size without first applying filtering.

Among the methods using shrinkage estimation, an advantage of *apeglm* is that it preserves true, large differences across conditions in the estimated LFCs. To demonstrate this, we calculated the average estimated LFCs for the methods that perform shrinkage *(apeglm, DESeq2, ashr, edgeR-PC5*), averaging over the 100 iterations. Comparing the average estimated LFCs to the reference LFCs demonstrates the extent of *bias* of the estimators, where it is expected that shrinkage estimators would have bias toward zero. We then constructed an MA plot, as typically used to visualize DE gene expression results, where we drew a *point* for the average estimated LFCs if it is within 0.5 units from the reference LFCs, or otherwise, we drew an *arrow* from the reference LFCs toward the average estimated LFCs (Supplementary Figure 7). All of the methods exhibit shrinkage of LFCs more than 0.5 for many genes with mean scaled counts less than 10, but *apeglm* preserved the most large LFCs for genes with larger mean scaled counts. *DESeq2* and *ashr* with *limma* input tended to shrink the LFCs by more than 0.5 for genes with mean expression levels greater than 10, including genes with absolute value of reference LFCs greater than 2, thus representing large differences across condition.

### Evaluation on simulation modeled on experimental data

We also checked whether *apeglm* provides accurate estimates of LFCs in simulated data modeled on experimental datasets. We generated the “true” LFCs from a mixture of zero-centered Normal distributions. The mean counts and Negative Binomial dispersion estimates were drawn from the joint distribution of the estimated parameters over the Pickrell et al. [20] and Bottomly et al. [17] datasets, as was performed in Love et al. [13]. We simulated 10,000 genes with a sample size of 5 vs 5, and repeated the whole simulation 10 times per experimental dataset. We also doubled the sample size to 10 vs 10 to see if the methods provided consistent relative performance at higher sample size. For the *Pickrell* dataset, which has higher within-group variation, we used a mixture of Normal distributions with standard deviations of 1, 2, 3 (with mixing proportions 90%, 5%, 5%, respectively). The *Bottomly* dataset has lower within-group variation, and so to make the simulation equally challenging, we used standard deviations of 0.25, 0.5,1 (90%, 5%, 5%). We constructed the simulation such that the expected count for all simulated samples was always greater than 10, to avoid overemphasizing the smallest count genes (this simulation choice does not advantage *apeglm*).

The simulation results for the *Pickrell* dataset (Figure 3) and the *Bottomly* dataset (Supplementary Figure 8) were mostly consistent with the previous result on the highly replicated yeast dataset. *limma, edgeR, edgeR-PC5*, and *apeglm* tended to have the lowest error when stratifying by true LFCs, although *limma* and *edgeR* had the lowest concordance when ranking genes by LFCs. The methods which do not shrink tended to produce large estimates for genes where the true LFCs are near 0 (Supplementary Figure 9 and 10). As in the yeast dataset, as the sample size increased, *apeglm* had lower error compared to *edgeR-PC5* for the largest LFCs. *DESeq2* had the highest error for the largest LFCs, as was expected. Unlike the results from the highly replicated yeast dataset, here *ashr* with both inputs had higher error for the middle range of LFCs. We note that we simulated *Negative Binomial* counts, and so the methods *apeglm, DESeq2* and *edgeR* which assume the Negative Binomial likelihood, are potentially at an advantage. *apeglm* struck a good balance for small and larger sample sizes, having consistently low error, and also high concordance in the CAT plot.

**Figure 3:**
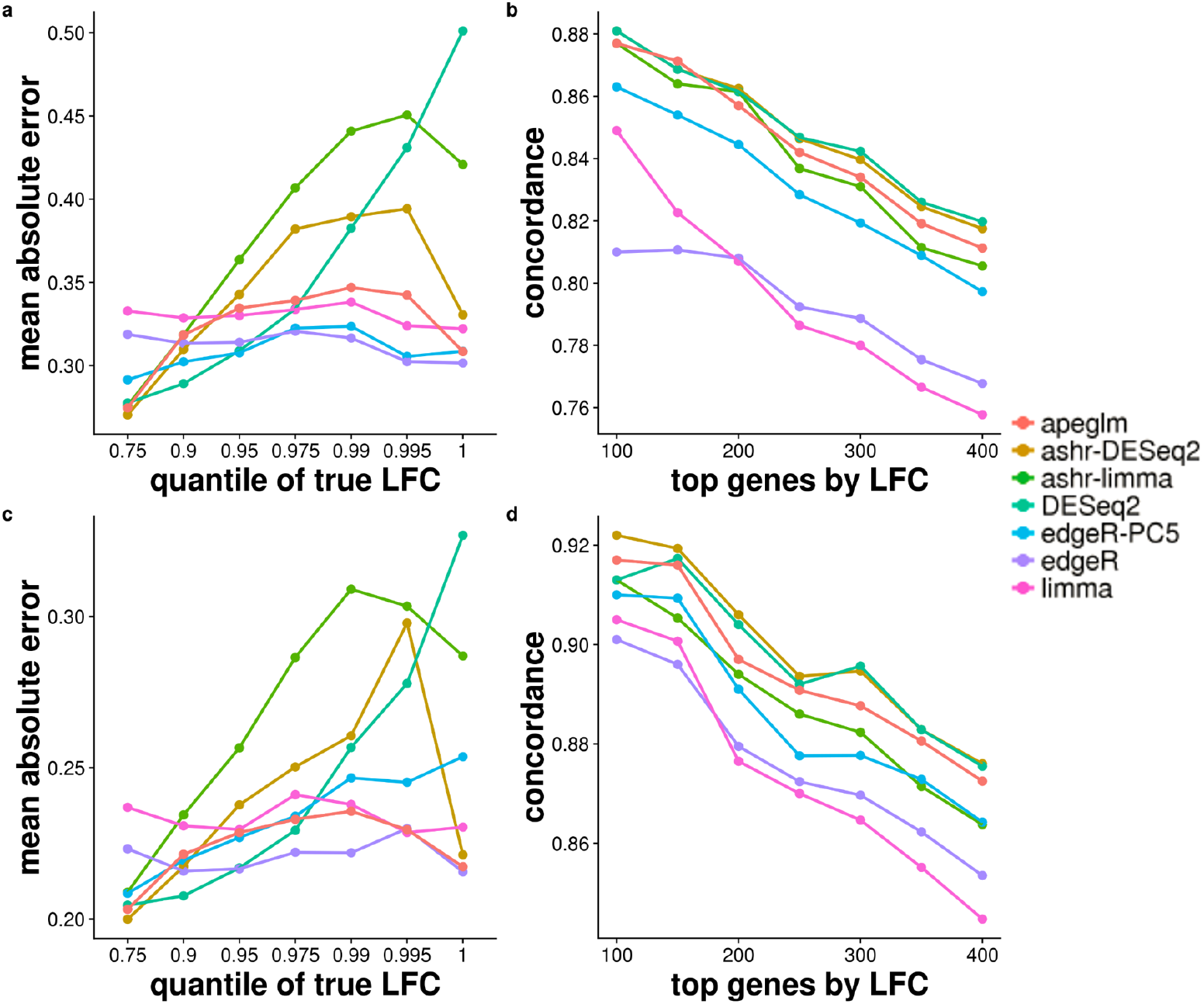
Simulation dataset (top row, 5 vs 5, and bottom row, 10 vs 10) modeled on estimated parameters from the Pickrell et al. [20] dataset. Each point represents the average over 10 repeated simulations.

The shrinkage estimators *apeglm, DESeq2*, and *ashr* tended to have low MAE across the range of counts (Supplementary Figure 11). *limma* and *edgeR* had high MAE for low counts.

The MAE for *edgeR-PC5* when binning genes by counts was low for the sample size of 5 vs 5, but higher when the sample size was increased to 10 vs 10.

Finally, we considered whether the methods which produce *s*-values (*ashr* and *apeglm*) were able to achieve their FSR bounds. We also generated *s*-values for *DESeq2* using the *DESeq2* posterior mode estimate and the associated uncertainty. We generated plots using the *iCOBRA* package [21], showing the number of genes at various achieved FSR values (Supplementary Figure 12). This analysis indicated that *apeglm* and *ashr* with *DESeq2* input tended to hit the target of 1% and 5% FSR, while *DESeq2* and *ashr* with *limma* input were just slightly above their nominal FSR. The *iCOBRA* data objects for four iterations of the simulation can be accessed at https://github.com/mikelove/apeglmPaper, and explored interactively using the *iCOBRA* Shiny app.

### Evaluation on cell line mixture experiment

We additionally evaluated the relative performance of *apeglm* using a cell line mixture RNA-seq dataset designed for benchmarking [22]. In this study, the investigators chose two cell lines from the same type of lung cancer, and grew the cell lines (NCI-H1975 and HCC827) as three

**Figure 4:**
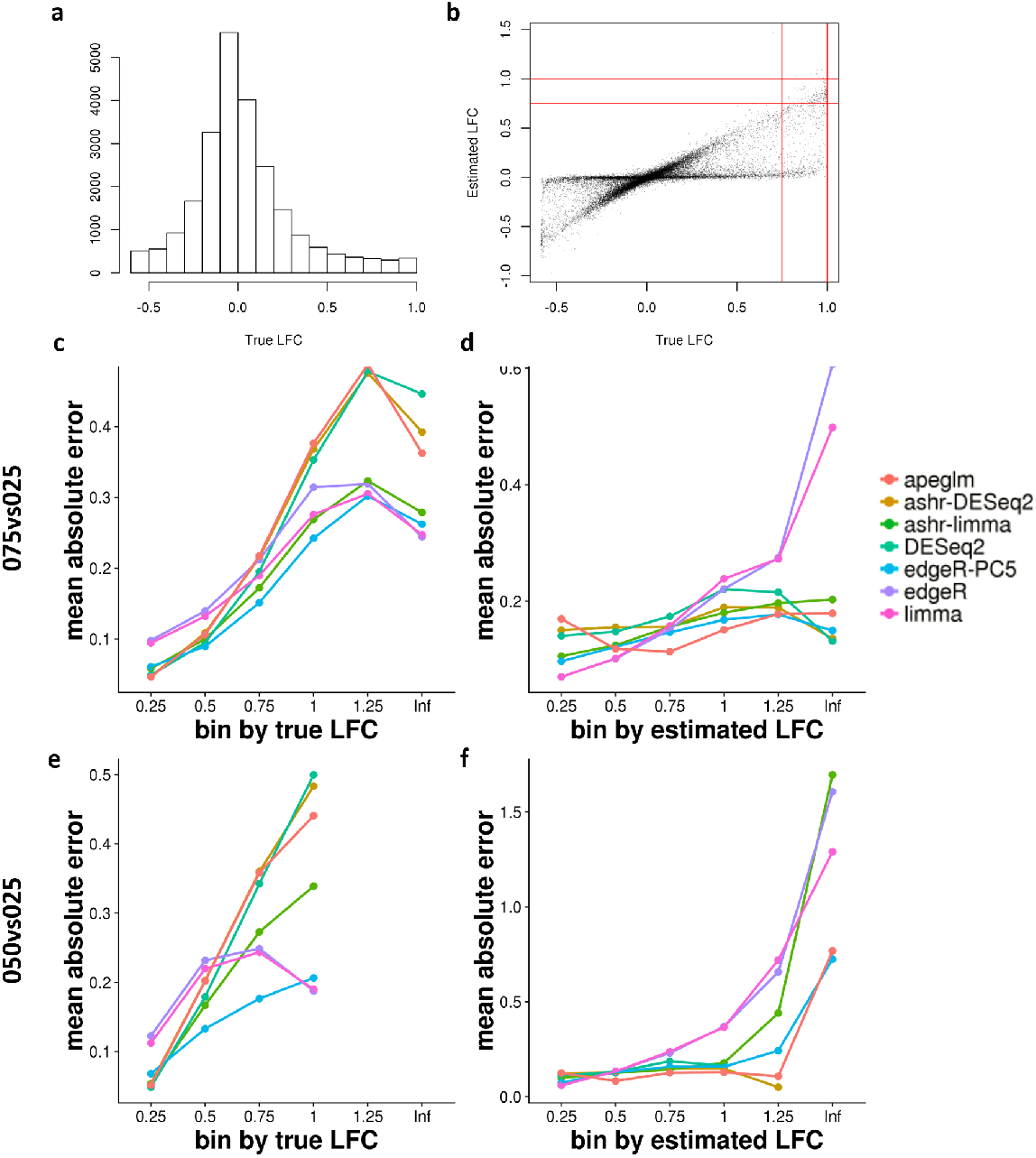
(a) The distribution of the true LFCs for comparison 050vs025, where the true LFCs is predicted with the fitted non-linear model. (b) Scatter plot of estimated LFCs from *apeglm* over true LFCs for comparison 050vs025. The vertical and horizontal lines indicate the two type of bins that were used for stratifying estimation error. (c) and (d) MAE plot binned by true LFCs and by estimated LFCs for comparison 075vs025. (e) and (f) MAE plot binned by true LFCs and by estimated LFCs for comparison 050vs025.

biological replicates, then mixed the RNA concentrations from each of these replicates at five pre-specified proportions (100%:0%, 75%:25%, 50%:50%, 25%:75%, 0%:100%). Following the notation of their paper, we use 100, 075, 050, 025 and 000 to represent the proportions. We used for evaluation the 15 normally processed samples prepared with Illumina’s TruSeq poly-A mRNA kit. We compared two groups of mixtures, each with three independent replicates: 075vs025 and 050vs025. We found the 100vs000 and 000vs100 mixtures were highly influenced by the 100 and 000 samples, which would be used both for estimation and for evaluation. We computed the estimation error as in Holik et al. [22] as the difference between the LFCs estimated by each method using two groups of samples and the LFCs predicted by a non-linear model fit to all 15 samples, using the `fitmixture` function in the *limma* package.

The distribution of true LFCs for the 075vs025 and 050vs025 are bounded by [log_2_ (1/3), log_2_ (3)] and [log_2_ (2/3), log_2_ (2)], respectively, and so instead of considering the top ranked genes, we considered two plots to assess the accuracy of LFC estimation: once binning by true LFCs and once binning by estimated LFCs (Figure 4). All shrinkage methods except *ashr* with *limma* input and *edgeR-PC5* had increased MAE for the highest LFCs when binning by true LFCs, although the shrinkage methods tended to perform well when binning by estimated LFCs. In these comparisons, *edgeR-PC5* tended to have consistently low error. We note the sample size for the cell line mixture experiment was 3 per group, and we expect the relative bias of the prior count approach to increase with sample size.

## Discussion

Here we compared various shrinkage estimators for LFCs in DE analysis of RNA-seq counts. RNA-seq experiments often have limited number of biological replicates in each condition group, typically in the range of 3 — 5. It is particularly difficult to estimate LFCs for genes with low counts or high coefficient of variation with such a small number of replicates. We examined methods for mitigating this problem of LFC estimation, and find that common filtering rules may lead to loss of DE genes. On the other hand, we found that existing methods for shrinking LFC estimates, such as *DESeq2*, may overly shrink those genes with very large LFCs, although the ranking was not greatly impaired. To reduce the shrinkage of large effect sizes that occurred using a *Normal* prior, we substituted an adaptive *Cauchy* prior, which has sufficient probability density in the tails of the distribution to allow for very large effects. The resulting estimator both reduced the variance associated with LFC estimates across the range from low to high counts, and also preserved true large LFCs.

## Conclusion

We have shown the utility in an adaptive, heavy-tailed prior for high-throughput experiments in which an effect size is estimated over tens of thousands of features. The results presented here have focused on the task of estimating the LFCs in RNA-seq experiments, using a Negative Binomial likelihood, but the software and methods are written in a general way, and in general, the use of the adaptive Cauchy prior may be adapted to other likelihoods and settings. The *apeglm* method accepts arbitrary likelihoods, as long as additional parameters are pre-specified, such as the dispersion. *apeglm* can therefore also be extended for use on other types of data, as long as it can be modeled by a GLM. For example, our method can be applied to allele-specific expression count data using a beta-binomial likelihood, as shown in the *apeglm* package vignette.

Providing low variance posterior mode effect sizes and their posterior standard deviation allows for various downstream uses, for example, plotting LFC estimates from two experiments against each other in a scatter plot, without having to make arbitrary filtering decisions that would have to apply to both datasets. In another context, the effect sizes of genetic variants across many different traits can be systematically correlated to one another to suggest potential relationships between the traits [23]. Such an analysis would benefit from shrunken estimates of effect size, to avoid hard filtering rules and to not have the correlations overly influenced by an imprecise outlier effect size estimate.

The computation of the approximate posterior provides useful aggregate statistics, such as the false sign rate and *s*-value proposed by Stephens [15], and the false-sign-or-small (FSOS) rate, which allows the user to define a range of effect sizes of biological significance. We note that, while the use of specific prior counts works well for providing point estimates of effect size for certain sample sizes and mean-variance relationships, it is difficult to choose a value that will work well for all datasets. For example, if one considers unique molecular identifiers (UMI) [24] and the counts produced following de-duplication in such an experiment, the information content of a low count can be much higher than in standard RNA-seq experiments without de-duplication, and so filtering rules and prior counts would need to be re-considered and manually adjusted for such a dataset. A Bayesian procedure for shrinkage of effect sizes, which takes statistical information into account, is desirable across different types of high-throughput datasets.

## Software versions

The following versions of software were used: REBayes 1.3, DESeq2 1.18.1, edgeR 3.18.1, limma 3.32.4, ashr 2.2-7, and apeglm 1.0.2.

## Declarations

### Ethics approval and consent to participate

Not applicable.

### Consent for publication

Not applicable.

### Availability of data and materials

The datasets analyzed during the current study are available in the ENA, GEO, or SRA repositories: Schurch et al. [19] https://www.ebi.ac.uk/ena/data/view/PRJEB5348, Holik et al. [22] https://www.ncbi.nlm.nih.gov/geo/query/acc.cgi?acc=GSE86337, Pickrell et al. https://trace.ncbi.nlm.nih.gov/Traces/sra/?study=SRP001540, Bottomly et al. [17] https://trace.ncbi.nlm.nih.gov/Traces/sra/?study=SRP004777.

*apeglm* is implemented as an R package and is available [25] as part of the Bioconductor project [26], at the following address: http://bioconductor.org/packages/apeglm. A single function apeglm is used to estimate the LFCs in the package, which takes a data matrix, a design matrix and a user-defined likelihood function as input. The function will return a list of estimated LFCs and corresponding posterior standard deviations, interval estimates, and arbitrary tail areas of the posterior. The *apeglm* package comes with a detailed vignette that demonstrates the functions in the package on a real RNA-seq dataset. The *apeglm* shrinkage estimator for RNA-seq can also be easily accessed from the *DESeq2* package, using the lfcShrink function. The R code used in this paper for evaluating methods is available at the following repository: https://github.com/mikelove/apeglmPaper

## Competing interests

The authors declare that they have no competing interests.

## Funding

MIL is supported by R01 HG009125, P01 CA142538, and P30 ES010126. JGI and AZ are supported by R01 GM070335 and P01 CA142538.

## Authors’ contributions

All authors developed the method and wrote the manuscript. AZ implemented the method and performed the analyses. All authors read and approved the final manuscript.

## Acknowledgments

The authors thank Wolfgang Huber and Cecile Le Sueur for helpful feedback on the software package.

## Methods

### Negative Binomial model for RNA-seq counts

We start with summarized measures of gene expression for the experiment, represented by a matrix of read or fragment counts. The rows of the matrix represents genes, (*g* = 1,…, *G*), and columns represent samples, (*i* = 1,…, *m*). Let *Y_gi_* denote the count of RNA-seq fragments assigned to gene *g* in sample *i*. We assume that *Y_gi_* follows a NB distribution with mean *μ_gi_*and dispersion *α_g_*, such that Var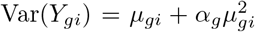. The mean *μ_gi_* is a product of a scaling factor *s_gi_* and a quantity *q_gi_* that is proportional to the expression level of the gene *g*. We follow the methods of Love et al. [13] to estimate *α_g_* and *s_gi_* sharing information across *G* genes, and consider estimates as fixed for the following. We fit a generalized linear model (GLM) to the count *Y_gi_* for gene *g* and sample *i*,

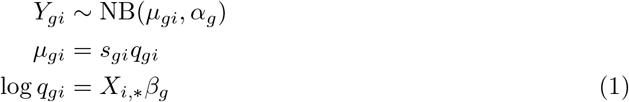

where *X* is the standard design matrix and *β_g_* is the vector of regression coefficients specific to gene *g*. Usually *X* has one intercept column, and columns for covariates, e.g., indicators of the experimental conditions other than the reference condition, continuous covariates, or interaction terms. We consider design matrices where the first element of *β_g_* is the intercept. For clarity, we partition the *β_g_* into *β_g_* = (*β_g0_*, *β_g1_*,…, *β_gK_*), where *β_g0_* is the intercept and *β_gk_*, *k*=1,…, K is for kth covariate. The scaling factor *s_gi_* accounts for the differences in library sizes, gene length [3] or sample-specific experimental biases [27] between samples, and is used as an offset in our model.

In the GLM, we use the logarithmic link function. In the *apeglm* software, the estimated coefficients and corresponding standard deviation estimates are reported on the same log scale. The *apeglm* method can be easily called from *DESeq2*’s lfcShrink function, which provides LFC estimates on the log_g_ scale. The *apeglm* method and software is generic for GLMs and can be used with other likelihoods. For example, it can be used for the Beta Binomial or zero-inflated Negative Binomial model, as long as estimates for the additional parameters, e.g. dispersion or the zero component parameters, are provided. An example of *apeglm* applied to Beta Binomial counts, as could be used to detect differential allele-specific expression, is provided in the software package vignette.

### Adaptive shrinkage estimator for *β_gk_*

We shrink coefficients representing differences between groups, continuous covariates, or interaction terms, but not the intercept. We propose a Cauchy distribution as the prior for the coefficients that the user wants to shrink. Therefore *β_gk_* in the model (1) has the prior

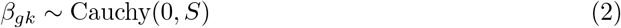

where the first parameter of the Cauchy gives the location and the second parameter is the scale, *S*. A similar default prior for coefficients associated with non-intercept covariates has been proposed by Gelman et al. [28] in the *bayesglm* R package, which uses a zero-centered Cauchy distribution with a scale of 2.5.

For setting the scale of the prior, we use the maximum likelihood estimates (MLE) 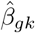 and their standard errors *e_gk_*. When making use of the set of MLEs for a coefficient, we shrink only a single coefficient at a time, and adapt the scale of the prior to the MLE by solving the following equation for *S*^2^ = *A* [29].

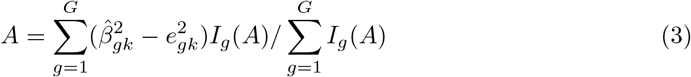

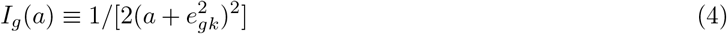

This equation is motivated by assuming that the MLE 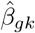 follows a Normal distribution around the true value *β_gk_* with variance 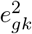, and that the *β_gk_* themselves follow a Normal distribution with mean zero. *A* is an empirical Bayes estimate of the variance of the *generating* Normal distribution, and 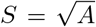 gives the scale. The equations above for estimating *A* are given by Efron and Morris [29], as a generalization of empirical Bayes estimators for the situation of many parameters each distributed with unequal variances. Equation 3 is solved for A using Brent’s line search implemented in R [30].

If the MLEs of the coefficients are not supplied, we use a scale *S* = 1 for all non-intercept coefficients. The unscaled posterior for *β_gk_* is the product of the prior density and the NB likelihood. We use the posterior mode, or *maximum a posteriori* (MAP), as the shrinkage estimator for the coefficient. The posterior mode is found using the L-BFGS algorithm [31] implemented in the *RcppNumerical* and *LBFGS++* libraries.

We derive the posterior distribution for *β_gk_* using the Laplace approximation: we estimate the covariance of the posterior distribution as the negative inverse of the Hessian matrix obtained from numeric optimization of the log posterior. We also attempted an alternate method for approximating the posterior by integrating the un-normalized posterior over a fine grid, but we found the Laplace approximation was consistently more accurate. Using the approximate posterior, we compute local FSR and credible intervals. Following Stephens [15], the local FSR is defined as the posterior probability that the posterior mode (MAP) is of the false sign, that is for gene *g*,

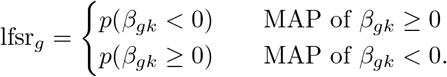

We also provide the local false-sign-or-smaller (FSOS) rate, relative to a given *θ* > 0 representing a biologically significant effect size,

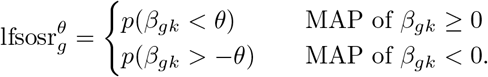

Analogous to the *q*-value [32], the *s*-value [15] provides a statistic for thresholding, in order to produce a gene list satisfying a certain bound in expectation. The *s*-value can be computed as

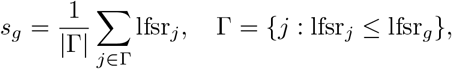

and likewise for the local FSOS rate. Other methods have suggested using the cumulative average or the cumulative maximum of posterior probabilities for defining the set of interesting features in high-throughput experiments include Leng et al. [10], Choi et al. [33], and Kall et al. [34].

### List of abbreviations

RNA-seq: RNA sequencing
ChIP-seq: chromatin immunoprecipitation followed by sequencing
LFC: logarithmic fold change
GLM: generalized linear model
DE: differential expression
FDR: false discovery rate
NB: negative binomial
MLE: maximum likelihood estimate
MAP: *maximum a posteriori*
CPM: counts per million
FSR: false sign rate
FSOS: false sign or smaller
SD: standard deviation
MAE: mean absolute error
CAT: concordance at the top
UMI: unique molecular identifier

---

**Supplementary Figure 1:**
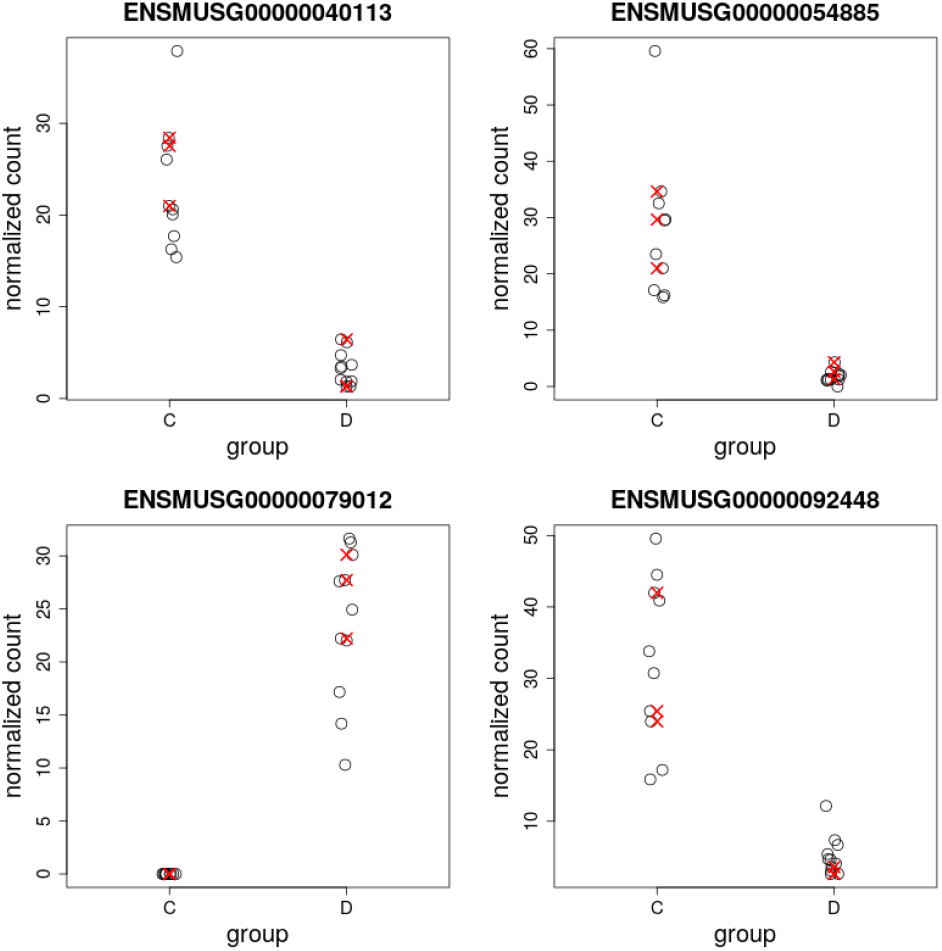
The scaled counts for four example genes. These four genes were filtered more than 50% of the time by a CPM filtering rule applied to 3 vs 3 samples, but were reported as DE by *DESeq2* in the full Bottomly et al. [17] dataset, with E(FDR) < 5%. The random subsets were balanced with respect to three batches and the full analysis controlled for batch in the design. The red X’s are examples of the scaled counts for one random subset, where this gene would be removed by the CPM rule, which requires ≥ 3 samples with CPM greater than the CPM value for a raw count of 10 for the least sequenced sample.

**Supplementary Figure 2:**
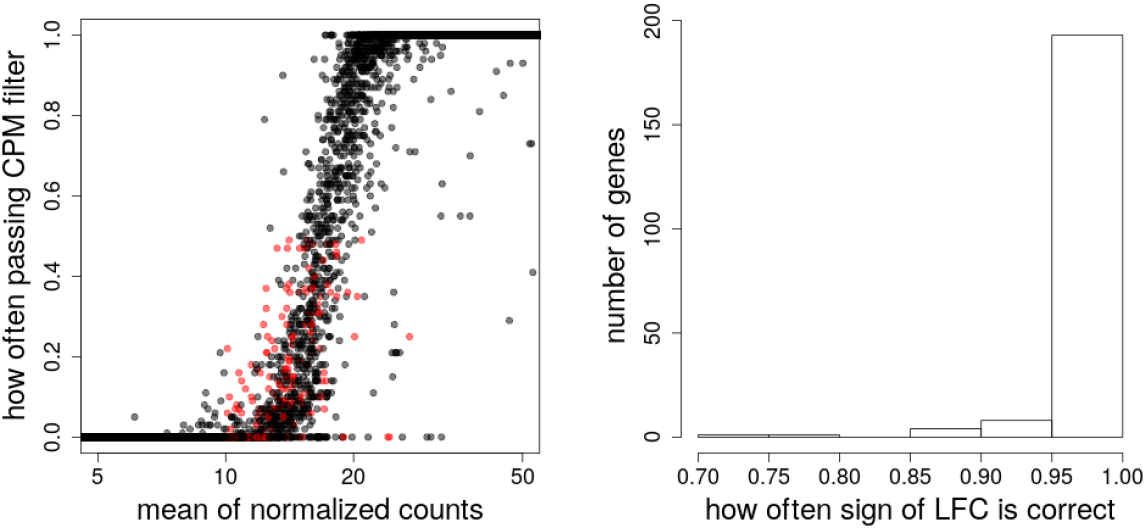
Details on 207 filtered genes. Shown in red are 207 genes (left) that were filtered more than 50% of the time by a CPM filtering rule in a 3 vs 3 random subset, which also have a mean of scaled counts in the full dataset greater than 10 and with E(FDR) < 5% on the full dataset. The histogram (right) indicates that these genes often had the correct sign of log fold change in random subsets (99% on average), indicating there was meaningful signal for these filtered genes.

**Supplementary Figure 3:**
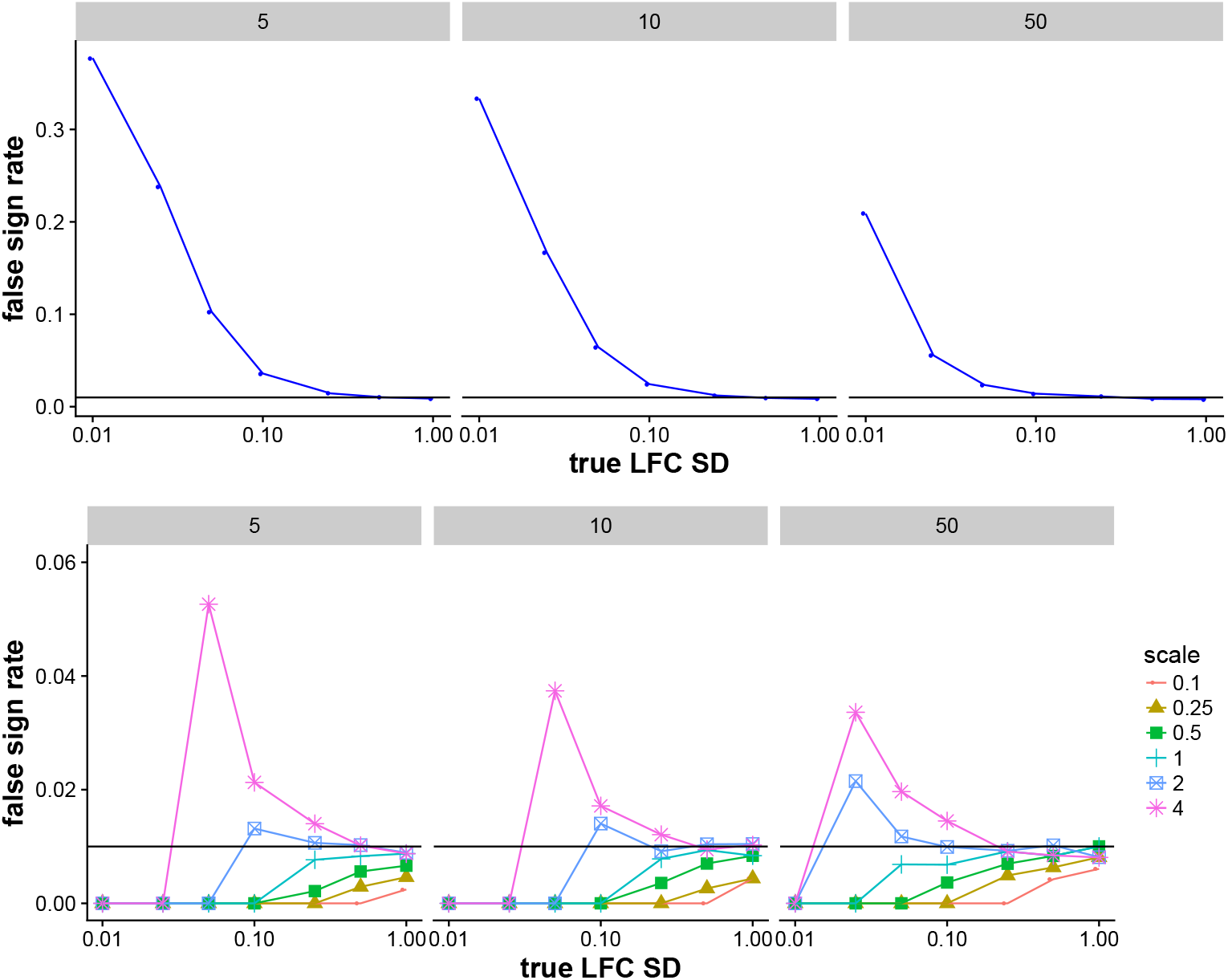
Achieved false sign rate on a simulation dataset. The top row represents using a fixed scaled (S=1) for the prior in *apeglm* while the bottom row represents scaling the prior to match the distribution of true LFCs, with various *multipliers* tested in the range [0.1, 4]. The three columns indicate different per-group sample sizes in {5,10, 50}. The points indicate the median over 100 iterations of the simulation. Here, the standard deviation (SD) of the distribution of true LFCs was provided as a *oracle* estimate to *apeglm* for setting the scale of the prior (*multiplier* * *true LFC SD*), whereas standard *apeglm* usage and for all other evaluations presented here the scale of the prior is estimated from the LFC MLE and their standard errors (Methods). The top row indicates that a fixed prior (S=1) did not control the FSR when the true LFC distribution was very narrow. The bottom row indicates that when the prior was 2 or 4 times the scale of the true distribution (blue square, pink asterisk), there was also some loss of FSR control, although to a lesser degree than using a fixed prior.

**Supplementary Figure 4:**
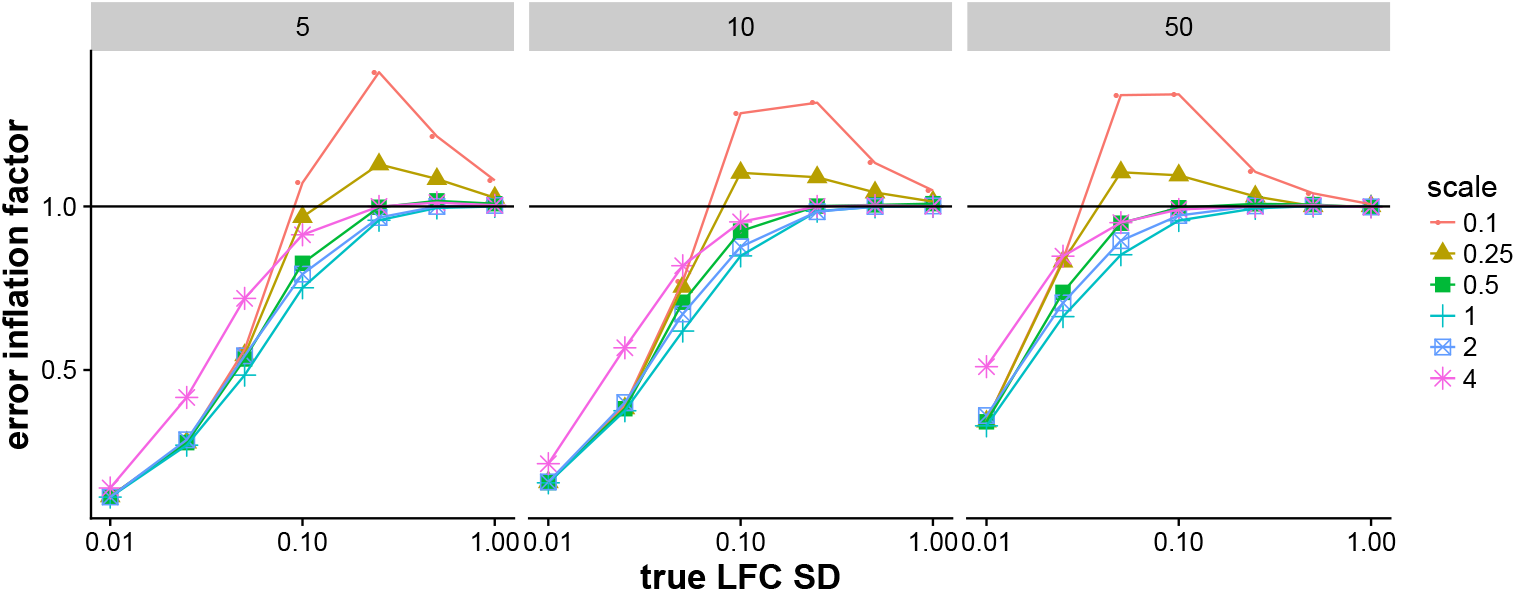
Error inflation factor for various scales of prior on simulated data. We define the error inflation factor as the ratio of mean absolute error (MAE) of the estimated LFC with an adaptive prior over the MAE of estimated LFC when using a fixed prior (S=1). For a middle range of true LFC SD, having a too narrow prior (red circles, yellow triangles) lead to an increase in MAE relative to a fixed prior. While in Supplementary Figure 3, scaling the prior to be *wider* than the true LFC SD caused a problem, here scaling the prior to be *narrower* than the true LFC lead to increased errors. Setting the scale of the prior equal to the scale of the true LFC SD struck a good balance (cyan plus sign). The following Supplementary Figure 5 shows what these errors looked like for individual genes.

**Supplementary Figure 5:**
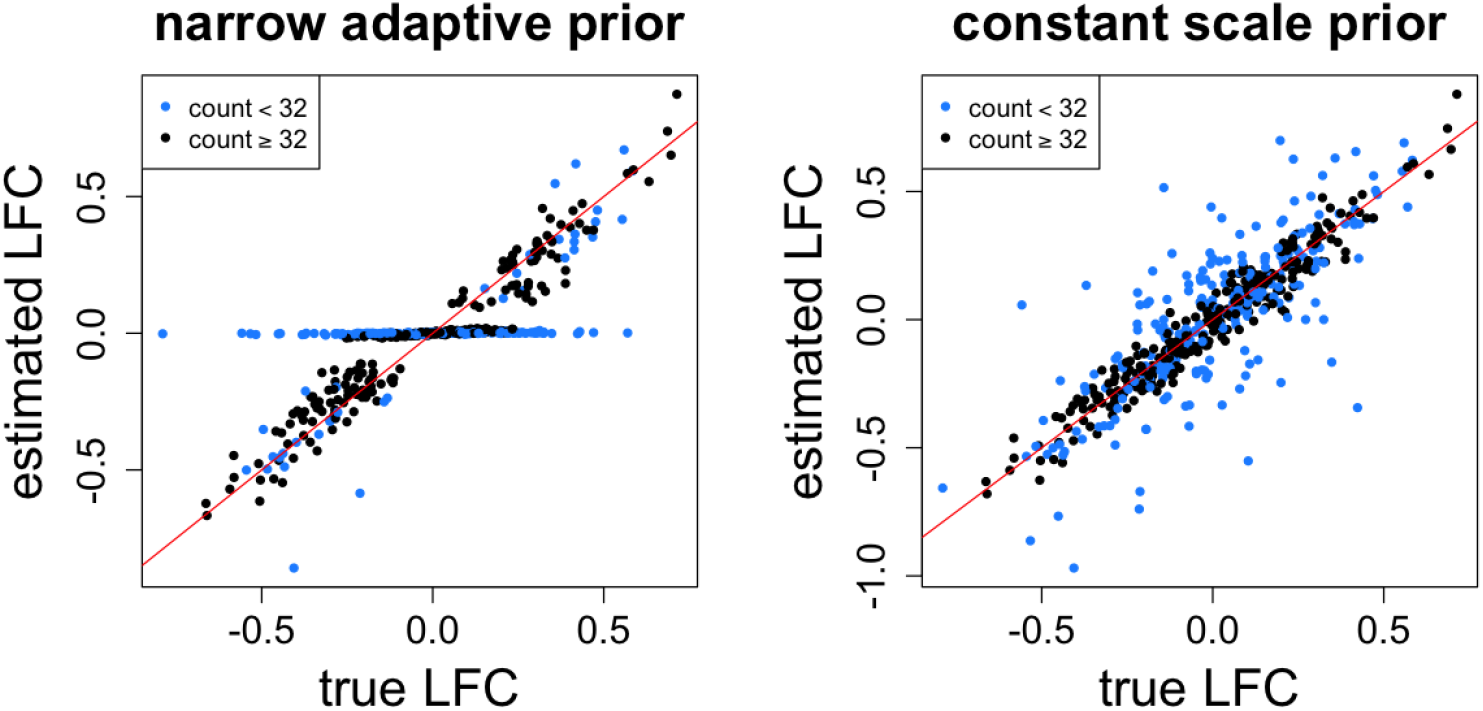
Examples of estimated LFC over true LFC for a too narrow prior and a fixed prior (S=1). Shown are the estimates for a single iteration of a 5 vs 5 sample comparison where the *true LFC SD* was 0.25 and the scale of the prior was 0.1 * *true LFC SD* = 0.025, i.e. using a *multiplier* of 0.1.

**Supplementary Figure 6:**
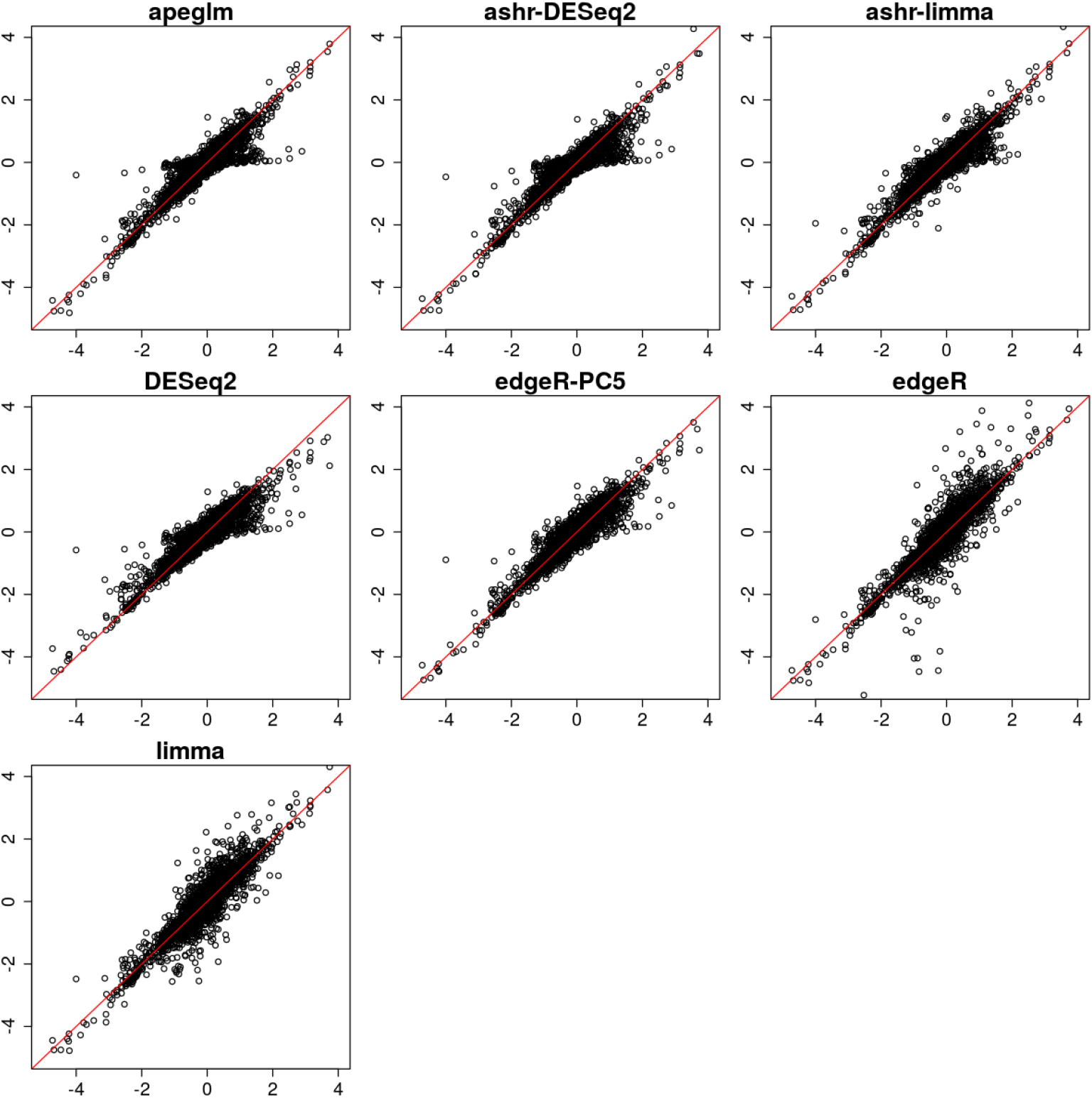
Estimated LFCs from a single iteration of a 3 vs 3 samples for all genes. The estimated LFC by seven different methods are plotted on y-axis against the reference LFC on x-axis. The red line denotes equality of estimated LFC and reference LFC. The bias for large effects for *DESeq2* can be observed, as well as the high variance for small effects for *edgeR* and *limma.*

**Supplementary Figure 7:**
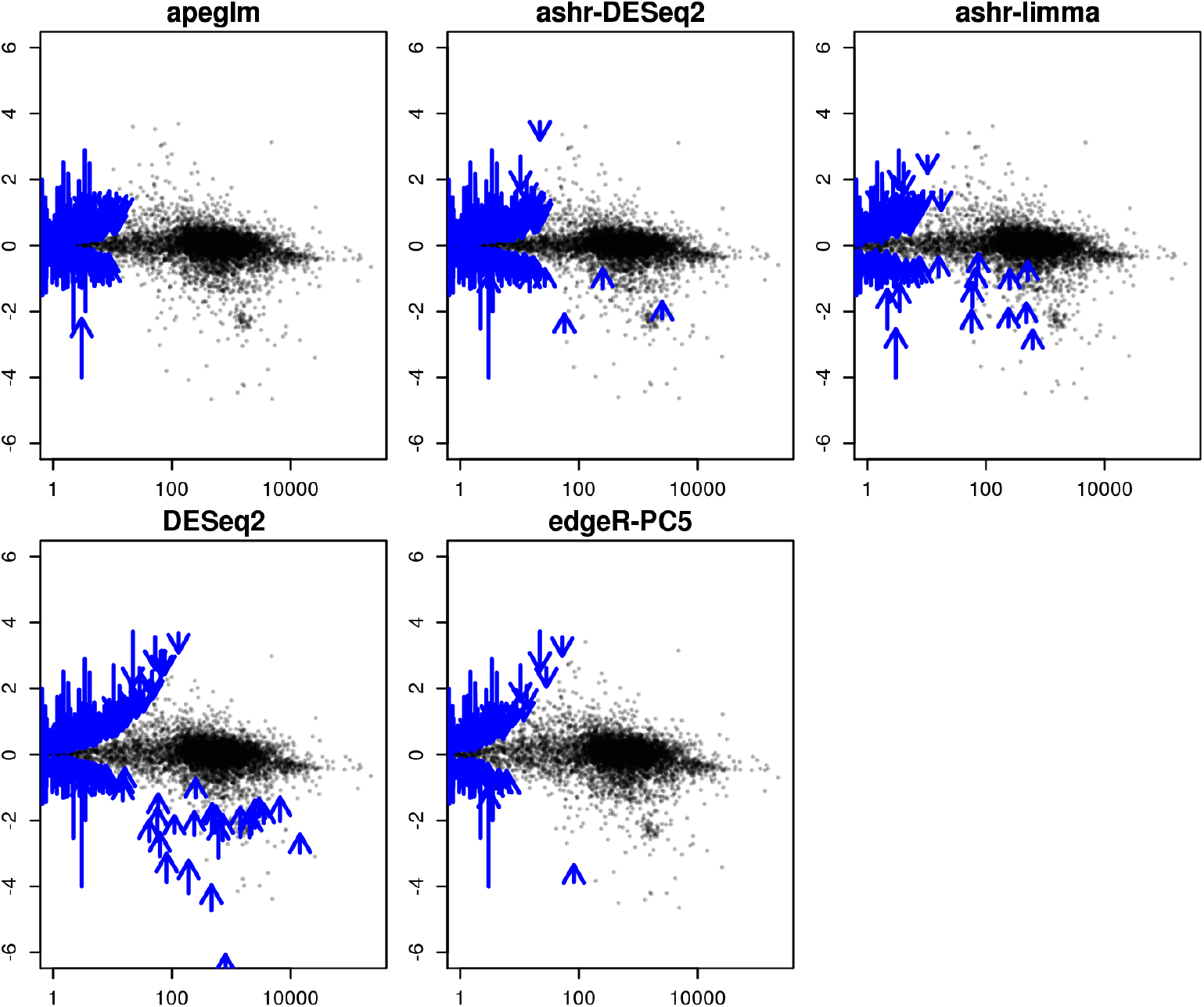
The MA plot of shrinkage estimates of LFC, averaging the estimates over 100 iterations for all genes from 3 vs 3 samples. The x-axis shows the mean of scaled counts from the full dataset. This MA plot visualizes the bias of shrinkage estimators across the range of mean signal. The points show the average estimated LFC when it was less than 0.5 units from the reference LFC. The arrows show when the difference between average estimated LFC and reference LFC was larger than 0.5. The tail of the arrow is the reference LFC and the head of the arrow is the average estimated LFC from the method.

**Supplementary Figure 8:**
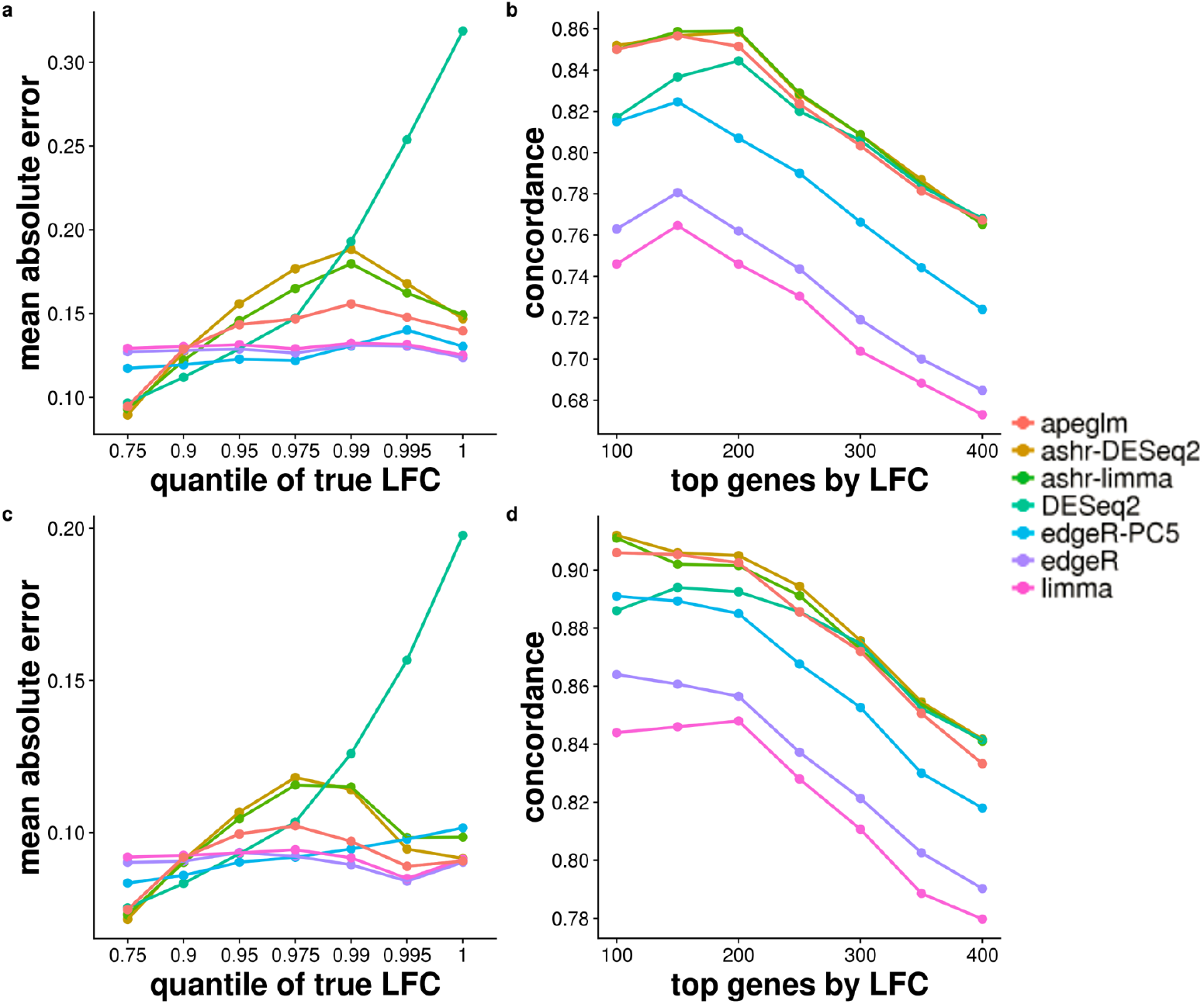
MAE plot and CAT plot of the simulation dataset (top row, 5 vs 5, and bottom row, 10 vs 10) modeled on estimated parameters from the Bottomly et al. [17] dataset. Each point represents the average over 10 repeated simulations.

**Supplementary Figure 9:**
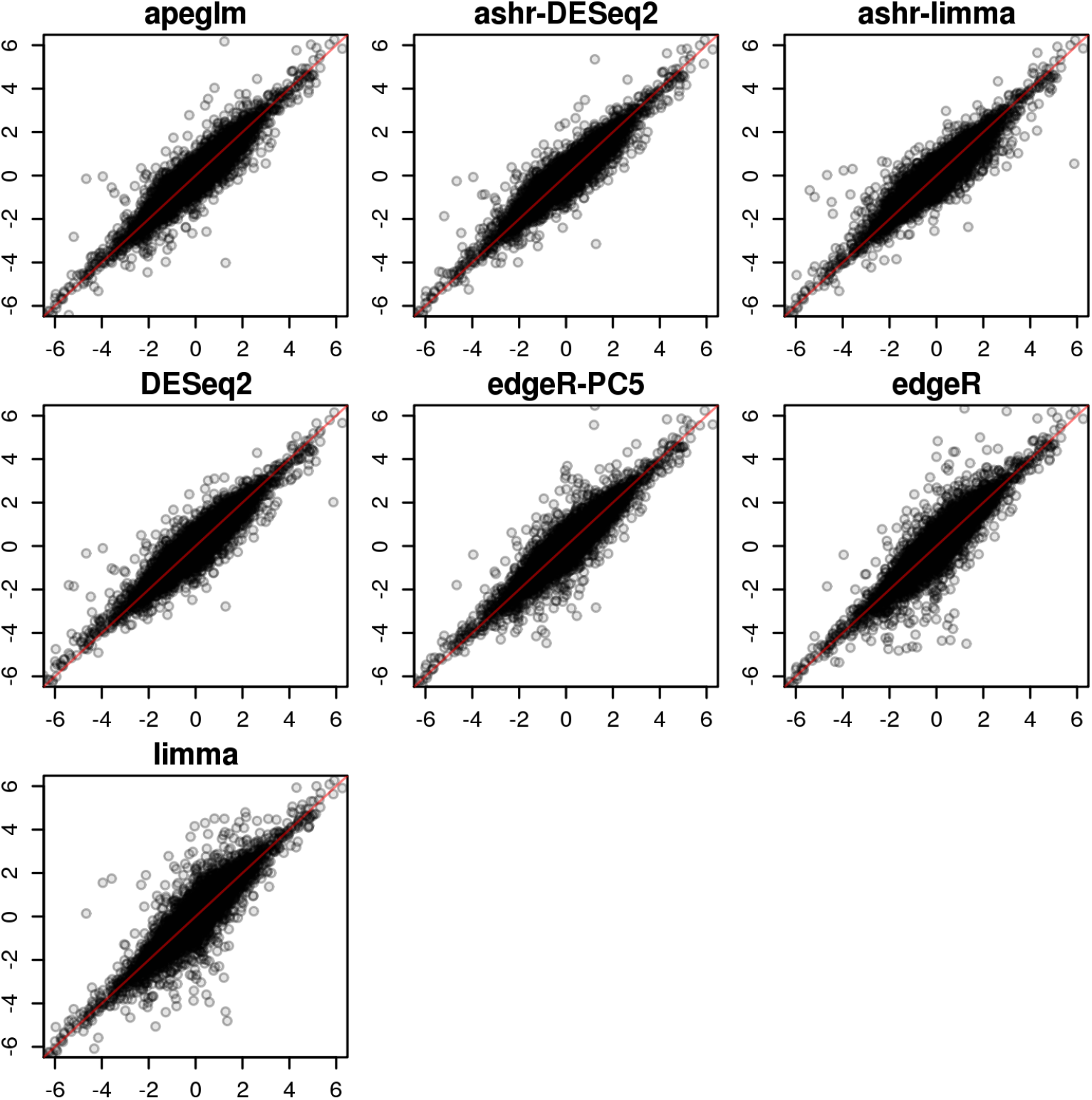
Scatterplot of estimated over true LFC for one iteration (5 vs 5) of the Pickrell et al. [20] dataset.

**Supplementary Figure 10:**
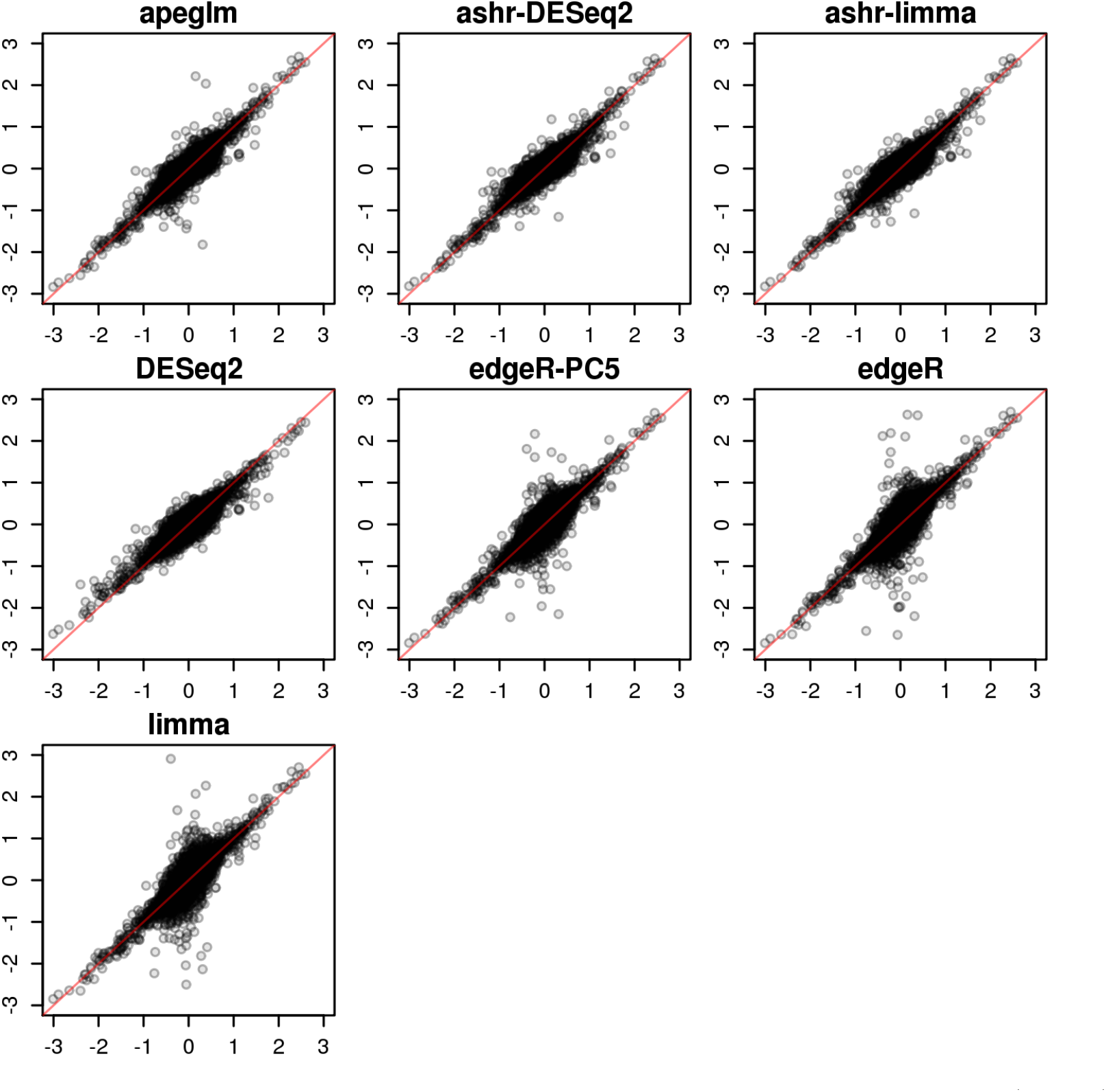
Scatterplot of estimated over true LFC for one iteration (5 vs 5) of the Bottomly et al. [17] dataset.

**Supplementary Figure 11:**
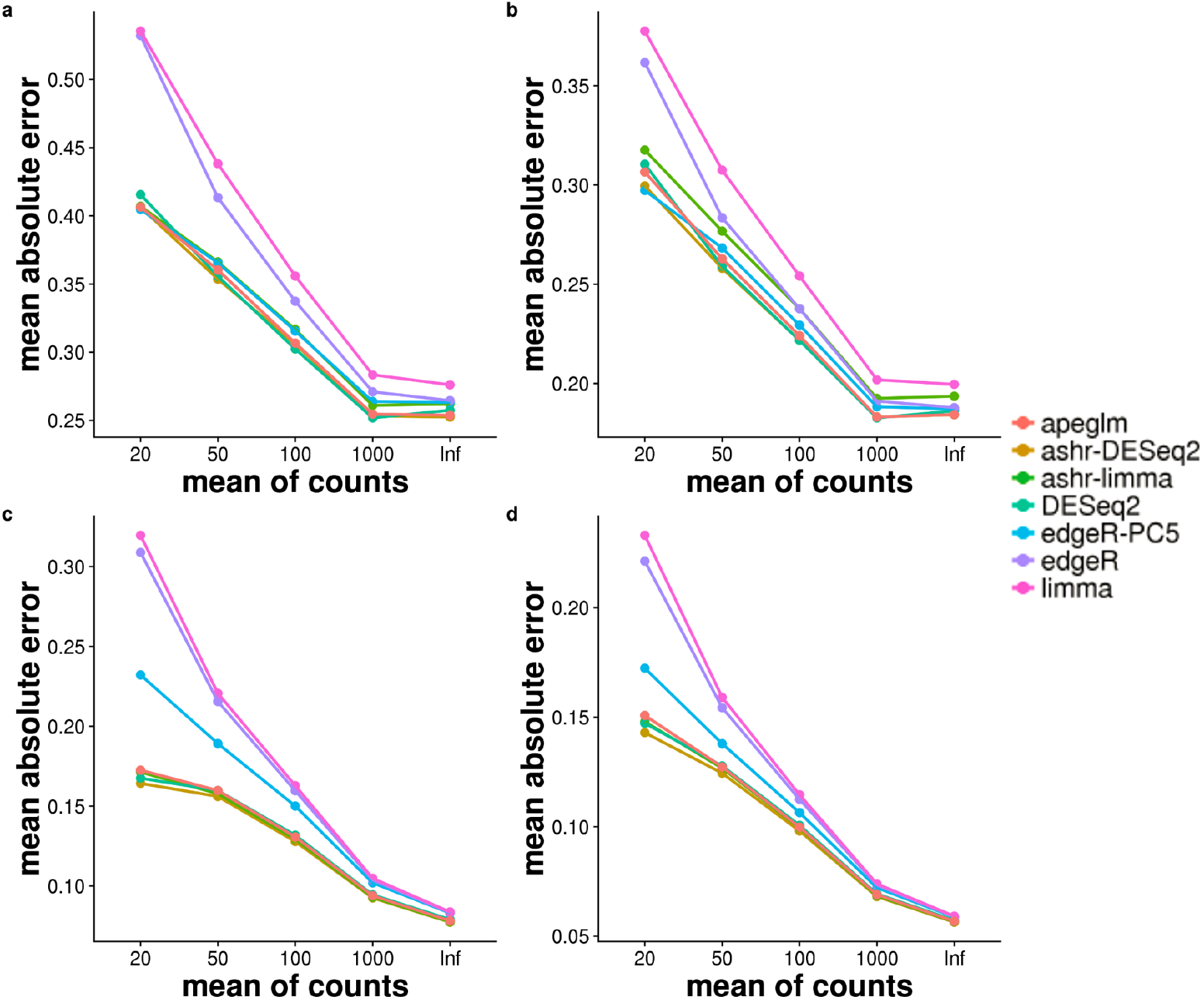
MAE plot binned by the mean scaled counts of simulation dataset (top row, 5 vs 5, and bottom row, 10 vs 10) modeled on estimated parameters from the Pickrell et al. [20] (a and c) Bottomly et al. [17] dataset (b and d). Each point represents the average over 10 repeated simulations.

**Supplementary Figure 12:**
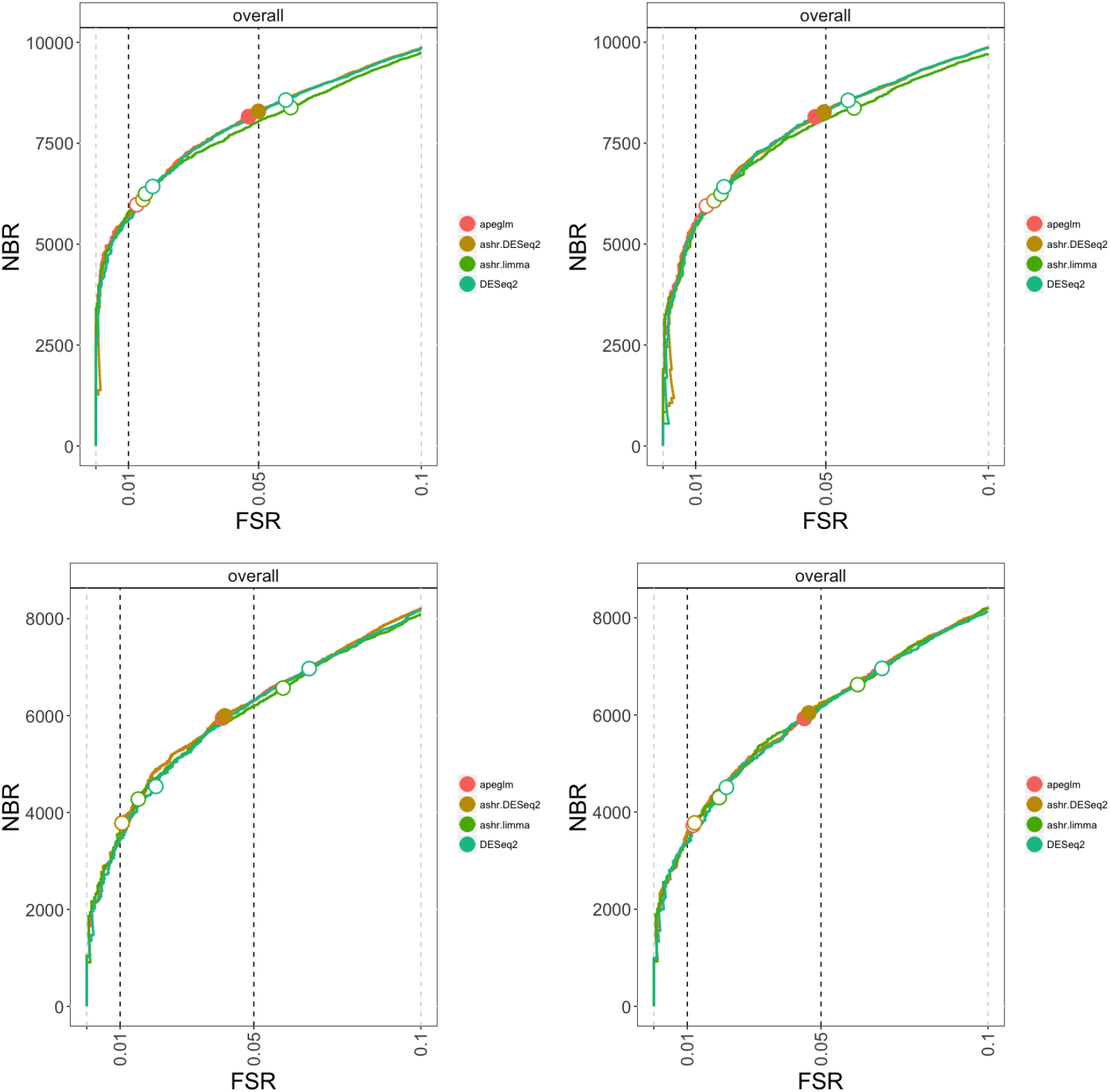
Number of genes (NBR) found at various achieved false sign rate (FSR). The top row shows two iterations of the simulated dataset modeled on the Pickrell et al. [20] data, the bottom row shows two iterations of the simulated dataset modeled on the Bottomly et al. [17] data, both with sample size of 5 vs 5. The plots are generated with the *iCOBRA* [21] Bioconductor package, with the two sets of circles indicating s-value cutoffs of 1% and 5%, and filled circles indicating that a method has achieved the nominal FSR bound from the s-value cutoff. The COBRAData objects for these four simulated datasets can be accessed at https://github.com/mikelove/apeglmPaper, where there are instructions on how to launch an interactive Shiny app for exploring the s-values and estimated LFCs across methods.

